# LAMPdecomposer: A generalisable framework for predicting and identifying loop-mediated isothermal amplification products using nanopore sequencing

**DOI:** 10.64898/2026.11.26.769529

**Authors:** Carlos A. dos Santos, Andrew McMahon, Nicole C Robb

## Abstract

Loop-mediated isothermal amplification (LAMP) is widely used for rapid molecular diagnostics, yet its complex concatemeric products remain poorly characterised, limiting assay validation and contamination detection. Here, we introduce LAMPdecomposer, a computational framework that combines a generalisable model of LAMP product formation with nanopore read decomposition to resolve individual amplification products. We predict 48 product sequences arising through four molecular mechanisms and validate their occurrence across 420,797 reads from 11 primer sets. Nearly all predicted products were recovered, revealing conserved patterns of product abundance and preferential amplification pathways. The library explained 52.2% of reads within its defined length range, with limited additional coverage from substantially longer reads. LAMPdecomposer also identified products originating from other primer sets at single-read resolution, revealing cross-contamination in positive samples and negative controls. By linking predicted amplification mechanisms to experimentally observed products, LAMPdecomposer provides a framework for investigating LAMP amplification and improving molecular diagnostic quality control.

## Main

Loop-mediated isothermal amplification (LAMP) has emerged as an important platform for rapid molecular diagnostics, particularly in point-of-care and resource-limited settings. Its development and diagnostic applications have expanded rapidly over the past decade, with further acceleration during the COVID-19 pandemic^1,2^. LAMP is a nucleic acid amplification technique widely used as an alternative to PCR^3^. Unlike PCR however, which requires precise thermal cycling, LAMP operates at a constant temperature of approximately 65 °C, which has enabled lab-on-a-chip and portable devices that bring molecular testing outside conventional laboratories^4–6^.

The features that make LAMP fast and sensitive also hinder its wider adoption. First, complex primer interactions make primer design laborious and empirical, because primer dimers can trigger exponential amplification even in the absence of target^7–9^. Second, the high product yield makes amplicon carryover a serious risk once reaction tubes are opened^10^. Third, LAMP generates a heterogeneous mixture of concatemeric products that usually require extensive processing before downstream analysis^11–13^. Together, these features make it hard to verify that the primers amplify only the intended target and to trace the source of false-positive amplification. Overcoming these problems requires an understanding of how LAMP products form, yet this process remains poorly characterised at the sequence level.

Kinetic and mathematical models have been proposed to explain the diversity of LAMP products^14,15^, but their validation has largely relied on gel electrophoresis. Electrophoresis can be used to separate the LAMP products by size but does not give information about the identity of the products. Interpretation is further complicated because the four to six LAMP primers can generate products through several alternative pathways, and structurally different products may be similar in size. Sequencing could in principle resolve these structures, but LAMP products are poorly suited to conventional short-read sequencing. Each product is a concatemer of near-identical target units repeated in alternating orientation and extending up to several kilobases, whereas short reads cover only a small part of a single molecule. Because the repeated units are almost identical, overlapping reads cannot be reliably reassembled into the original molecules, and the order, number and orientation of units are lost. Consequently, existing sequencing-based LAMP assays fragment or PCR-amplify the products and use the reads only to confirm the presence of the target sequence^16,17^, rather than to determine product structure.

Here, we developed a pipeline to resolve LAMP products directly, by modelling a reaction product tree comprising 48 sequences expected from specific amplification. We validated the tree using a published library of Oxford Nanopore-sequenced LAMP products, in which each read corresponds to a single amplified molecule, allowing the relative abundance of each product to be estimated. We showed that this tree accounts for the majority of LAMP products, describe how these products form, and generalise our findings into a framework applicable to any LAMP target. We also present LAMPdecomposer, a Python-based tool that uses the StringDecomposer algorithm^18^ to decompose Oxford Nanopore reads from any LAMP target, with sufficient sensitivity to detect a single contaminating read from another target.

## Results

### Creating a base library of expected amplification products

To characterise the diversity of LAMP amplification products, we sought to establish a generalisable model capable of predicting the sequences generated during amplification, independently of the target or primer set. We hypothesised that, despite the apparent complexity of LAMP products, their diversity could be explained by a limited number of amplification mechanisms acting on a defined set of intermediate structures.

Amplification starts when the inner primers (FIP/BIP) invade the target double-stranded DNA (dsDNA), a process facilitated by the loop (LF/LB) and outer (F3/B3) primers (Figure 1A)^19^. The following two rounds of amplification generate an amplification seed containing the inner primer sequences at both ends. This molecule can exist either as dsDNA or as the dumbbell structure characteristic of LAMP. Starting from this initial seed, we identified four mechanisms capable of generating the products in our model: (i) extension of single-stranded DNA (ssDNA) into blunt-ended products, (ii) recycling of amplification seeds, (iii) extension from a self-folded 3′ end and (iv) annealing and extension of two products with complementary 3′ regions (Figure 1Bi-iv).

**Figure 1.**
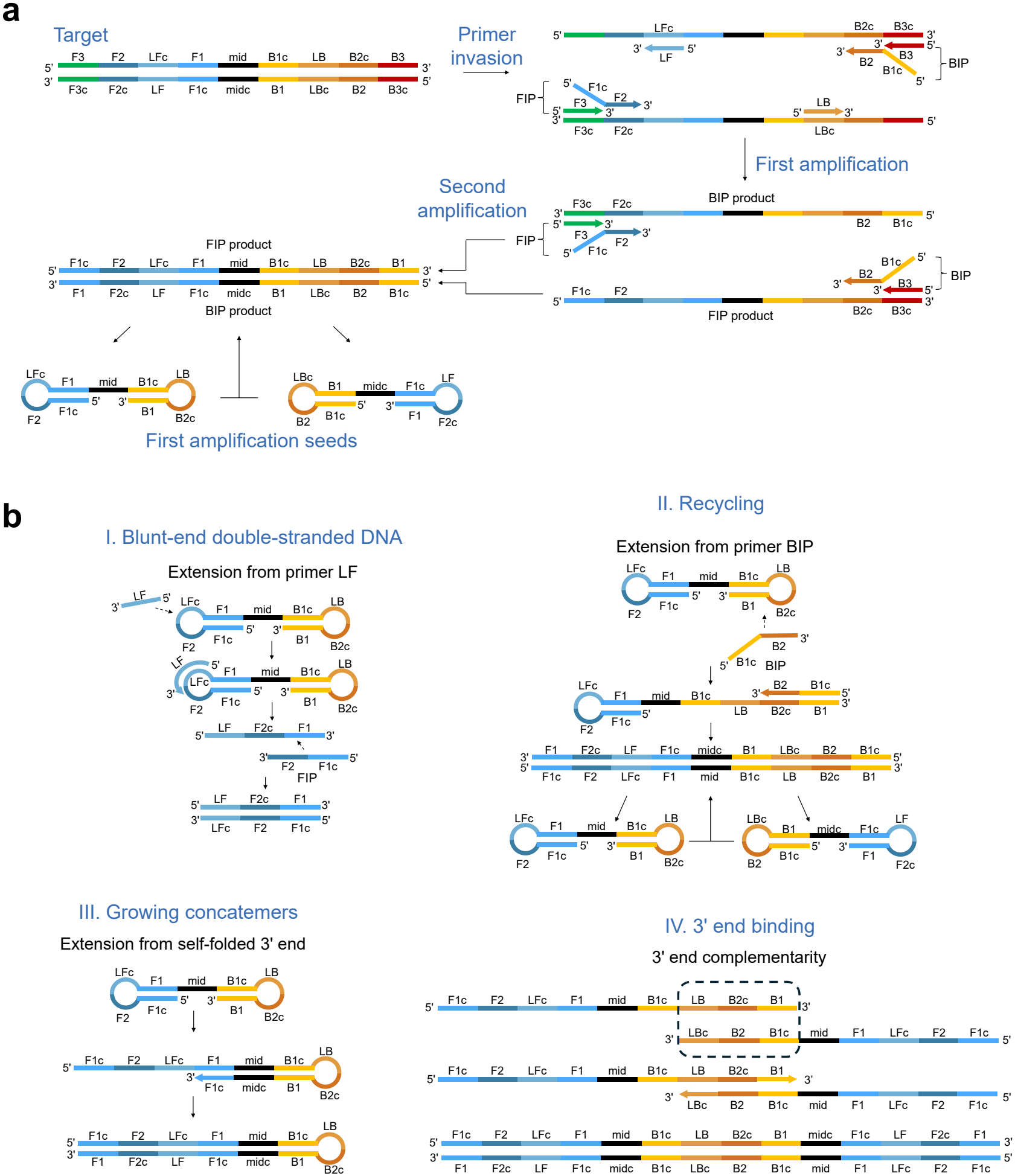
Mechanisms of LAMP product formation. **a**. Generation of the first amplification seed. In the first amplification, the inner primers (FIP/BIP) invade the target dsDNA and are extended, and the outer primers (F3/B3) displace the newly synthesised strands. In the second amplification, the inner primers bind these products, generating an amplification seed with inner primer sequences at both ends. The seed can fold into two dumbbell structures. **b**. The four mechanisms that explain all products in the base library. (i) Extension of ssDNA into blunt-end dsDNA. (ii) Recycling of amplification seeds. (iii) Growing concatemers. (iv) 3′ end binding. Colours indicate F side regions (blue), the region between F1 and B1c (mid, black) and B side regions (orange). A suffix c denotes the reverse complementary sequence.

The first mechanism accounts for the formation of stable, blunt-ended dsDNA molecules. For example, the LF primer can extend the amplification seed to generate a product which is subsequently extended by the FIP primer, creating a duplex (Figure 1Bi). Although these products cannot fold into longer concatemers, they can undergo repeated primer invasion and copying. The second mechanism describes the recycling of amplification seeds, whereby extension from the inner primers regenerate the initial seed structures, allowing further rounds of amplification (Figure 1Bii). The third mechanism accounts for concatemer growth through self-priming: sequences ending in the reverse complement of FIP or BIP can fold back to form a hairpin, enabling extension from their own 3′ ends and generating products longer than the original molecules (Figure 1Biii). The fourth mechanism provides an alternative pathway for product formation through the annealing of molecules with complementary 3′ ends, for example, products that terminate in the complementary regions 5′…LB-B2c-B1 3′ and 5′…B1c-B2-LBc 3′, respectively. Annealing of these regions allows extension of both strands, regenerating a product from pathway iii and generating a new complementary product (Figure 1Biv). Thus, individual products may arise through more than one amplification pathway, further contributing to the complexity of LAMP reactions.

Based on these mechanisms, we constructed a base library comprising 24 LAMP products and their reverse complements (48 sequences in total) (Figure 2A & Supplementary Figure 1). The library assumes that, although LAMP amplification generates numerous displaced ssDNA molecules, these can ultimately form stable dsDNA products. Accordingly, displaced strands that fold into hairpins are treated as intermediates that can subsequently be extended into complete dsDNA molecules. These products can, in turn, undergo further primer invasion and extension, releasing additional ssDNA strands and sustaining amplification. This approach allowed us to represent the diversity of amplification products as a defined set of sequences connected through specific molecular mechanisms (Figure 2B). The map shows only the main route to each molecule and omits mechanism (iv) whenever another pathway explains it. All amplification pathways can be explored interactively in Supplementary File 1. Importantly, because the model is defined by the arrangement of primer-binding regions rather than target-specific nucleotide sequences, it can be adapted to different LAMP assays. This provides a basis for testing whether the predicted products are present in experimental reactions and determining how much of the observed product diversity can be explained by the model.

**Figure 2.**
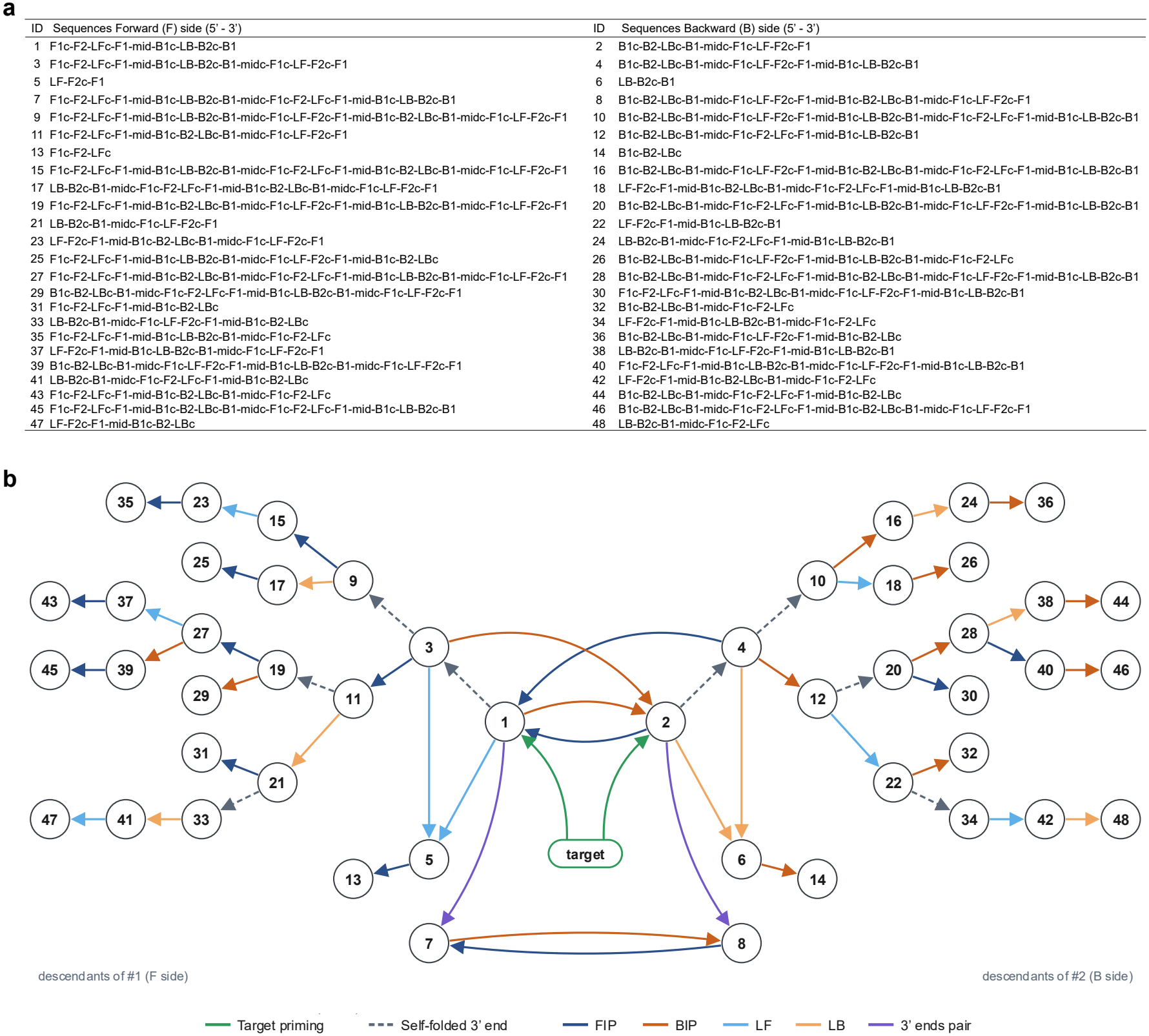
The LAMP base library and amplification product tree. **a**. The 48 sequences of the base library, written 5′ to 3′ as strings of primer target regions. Odd IDs are Forward side (F side) sequences and even IDs are Backwards side (B side) sequences. **b**. Product tree showing how each sequence is generated from the target. Nodes are sequence IDs from a. Arrows indicate the mechanism of formation. The first target priming is represented as a single green ellipse, extension from a self-folded 3′ end is shown in grey dashed lines, extension from FIP is represented in dark blue, BIP in brown, LF in light blue, LB in light orange, and 3′ end pairing in purple. Descendants of #1 (F side) are shown on the left and descendants of #2 (B side) on the right. For clarity, only the main route to each molecule is shown, and 3′ end pairing is omitted whenever another pathway explains the molecule.

### The base library captures the primary LAMP amplification products

To determine whether the predicted amplification products occur in experimental LAMP reactions, we required a method capable of resolving the sequence composition of individual concatemeric molecules. Unlike conventional short-read sequencing, nanopore sequencing can generate reads spanning entire LAMP products, allowing their constituent regions to be identified and their structures compared with the predicted library. However, the repetitive and variable arrangement of these regions presents a challenge for conventional sequence analysis.

The StringDecomposer algorithm was developed to decompose reads from the repetitive regions of centromeres into their constituent monomers, translating them from the nucleotide alphabet into strings of monomers^18^. We hypothesised that the same software could resolve LAMP concatemeric structures if each primer target region were treated as a monomer. We therefore developed LAMPdecomposer, a tool that adapts the base library to each primer set, then uses StringDecomposer to assign monomers along each read and uses a post-processing step to assemble these assignments into a monomer pattern (Figure 3A). Each pattern is then matched against the version of the base library built for each primer set. LAMPdecomposer reports the proportion of reads explained by the library and the percentage of predicted products found in each sample.

**Figure 3.**
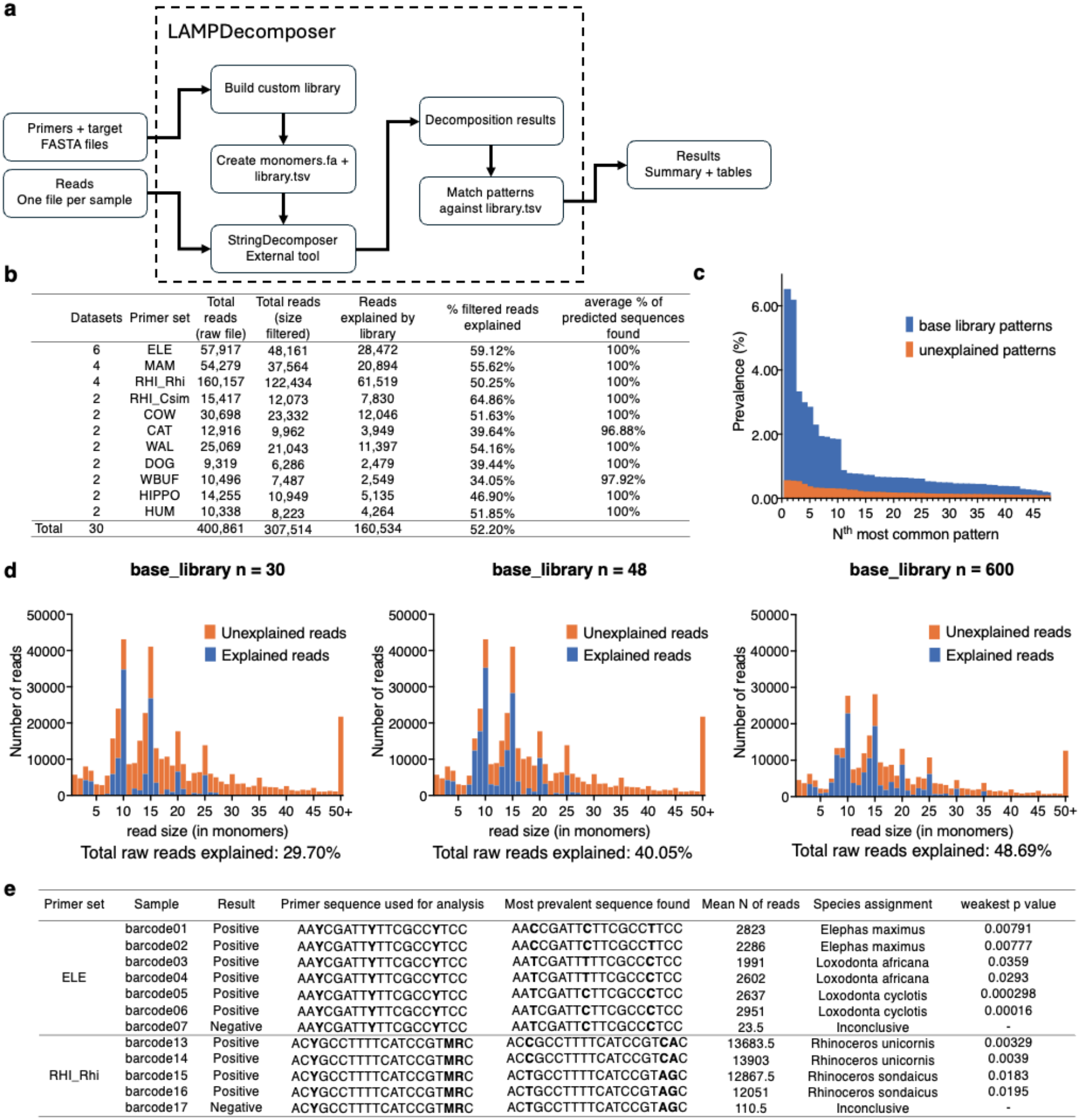
The base library explains the most common LAMP amplification products. **a**. LAMPdecomposer workflow. From the primer and target FASTA files, LAMPdecomposer builds a custom library for each primer set (monomers.fa and library.tsv). Reads, provided as one file per sample, are decomposed into primer target monomers with StringDecomposer. The decomposition output is then assembled into monomer patterns and matched against the library, producing summary tables for each sample. **b**. Results for the 30 positive amplifications (400,861 reads) across 11 primer sets. Filtered reads exclude reads shorter than the shortest or longer than the longest library sequence, measured in monomers. The total percentage of filtered reads explained is pooled across all datasets. The final column gives the percentage of the 48 predicted sequences found, averaged across datasets for each primer set. **c**. Prevalence of the 48 most common patterns, averaged across positive samples. Base library patterns (blue) and unexplained patterns (orange) are ranked separately. **d**. Read size distribution for libraries of 30, 48 and 600 sequences. The 600 pattern library was built from the most common patterns observed in the ELE and MAM primer sets, so this library was evaluated on the remaining nine primer sets. Bars show the total reads explained by the library (blue) and unexplained (orange). The y axis shows the total number of reads per size bin across all raw samples, and the final bin pools reads of 50 or more monomers. The percentage of total raw reads explained is shown below each panel. **e**. Species assignment with SNPdetector.py for primer sets targeting more than one species (ELE and RHI_Rhi). Reads were searched for every version of the degenerate LB primer, and its reverse complement. A species was assigned when the most abundant version was significantly more abundant than every other version (one-sided two-sample t-test, p < 0.05). The p value shown is the largest of these comparisons. Barcodes 07 and 17 are no-template controls. The raw data is from Yugovich et al.^20^.

To assess how well the base library explains the diversity of LAMP amplification products, and how this depends on library size, we analysed a published dataset of 41 raw FASTQ files generated with 11 primer sets, comprising 420,797 reads^20^. Of these, 400,861 reads came from 30 positive amplifications and the remaining 19,936 from 11 negative samples. The primer sets used in this dataset target species-specific single nucleotide polymorphisms (SNPs) in the mitochondrial 16S rRNA, 12S rRNA and cytochrome b (CytB) genes of species relevant to the illegal wildlife trade: elephant (*Elephas maximus, Loxodonta africana* and *Loxodonta cyclotis*; ELE), mammoth (*Mammuthus primigenius* and *Mammuthus columbi*; MAM), rhinoceros (*Rhinoceros unicornis* and *Rhinoceros sondaicus*, RHI_Rhi; *Ceratotherium simum simum*, RHI_Csim), cow (*Bos taurus*; COW), cat (*Felis catus*; CAT), walrus (*Odobenus rosmarus*; WAL), dog (*Canis lupus familiaris*; DOG), water buffalo (*Bubalus bubalis*; WBUF), hippopotamus (*Hippopotamus amphibius*; HIPPO) and human (*Homo sapiens*; HUM). Our aim was not to explain every read, which would in principle be possible with a sufficiently large base library, but to identify the smallest set of sequences that explains the observed reads satisfactorily.

We first assessed whether the predicted products were consistently detected across primer sets, before determining what proportion of sequencing reads could be assigned to the library. Nearly all predicted products were detected in every primer set. On average, all 48 sequences of the base library were found in nine of the 11 primer sets, and 96.88% and 97.92% were found in CAT and WBUF, respectively (Figure 3B). We then excluded reads that the library could not explain by definition, namely those shorter than the shortest or longer than the longest library sequence, measured in monomers. Overall, the 48 sequences explained 52.20% of the remaining 307,514 reads, ranging from 34.05% for WBUF to 64.86% for RHI_Csim (Figure 3B).

A small number of products dominated the reaction. The most prevalent product, counting both its forward and reverse complement sequences, represented 12.7% of the size filtered products on average, and each subsequent product explained progressively less (Figure 3C). Unexplained patterns followed the same trend. The most common were only slightly more prevalent than the least common library sequences, and each additional unexplained pattern accounted for an ever-smaller share of the reads (Figure 3C).

We therefore tested whether a substantially larger library could meaningfully improve read classification by comparing three libraries: the first 30 sequences of the amplification product tree, all 48 sequences in the tree, and the 600 most common patterns observed in two primer sets (ELE and MAM). Because the 600-pattern library was built from observed rather than predicted sequences in ELE and MAM, the 600-sequence library was evaluated on the remaining nine primer sets. Increasing the library from 30 to 48 sequences raised the proportion of explained reads from 29.70% to 40.05%. Expanding it to 600 patterns, more than 12 times as many sequences, explained 48.69%, an increase of only 8.64 percentage points (Figure 3D). Even when evaluating the 600 most common patterns, the ones in the 50+ monomer bin remained largely unexplained.

Together, these results show that a relatively small, mechanistically derived library captures nearly all predicted product types across diverse LAMP assays, while explaining a substantial proportion of sequencing reads. The limited improvement achieved by expanding the library suggests that much of the remaining sequence diversity comprises low-abundance products or structures beyond the length range of the predicted library. Importantly, the consistent recovery of the predicted products across primer sets supports their use as sequence-level markers of specific LAMP amplification, providing a basis for identifying products originating from other targets.

### Identifying products amplified from closely related species

Because LAMPdecomposer assigns reads to a primer set by matching entire monomer regions, it tolerates single-nucleotide differences between reads. This makes read assignment robust to sequencing errors, but it also prevents LAMPdecomposer from distinguishing closely related species when one primer set targets several of them. We therefore developed a companion script, SNPdetector.py, that counts the reads carrying each species-diagnostic SNP. Two primer sets targeted more than one species: ELE (Elephas maximus, Loxodonta africana and Loxodonta cyclotis) and RHI_Rhi (Rhinoceros unicornis and Rhinoceros sondaicus). In both sets, the LB primer contains three degenerate positions at the nucleotides that differ between these species, so the primer version that matches the template should dominate the amplicons. For each LB version, SNPdetector counted the reads containing its sequence and, separately, its reverse complement. Because LAMP concatemers contain alternating sense and antisense copies of the target, these two counts should be similar, so we treated them as two measurements of the same version. The most abundant version was compared with every other version (one-sided two-sample t-test), and a sample was assigned to a species only when that version was significantly more abundant than all others (p < 0.05). All ten samples could be assigned to one of the tested species (p = 0.00016 to 0.0359). Barcodes 01 and 02 were assigned to *E. maximus*, 03 and 04 to *L. africana*, 05 and 06 to *L. cyclotis*, 13 and 14 to *R. unicornis*, and 15 and 16 to *R. sondaicus* (Figure 3e). As expected, the no-template controls (barcodes 07 and 17) gave no call. These results corroborate the findings of the original paper by Yugovich et al.^20^

### LAMPdecomposer is sensitive enough to detect single-read cross-contamination across samples

By running LAMPdecomposer on each sample with each primer set as monomers, we can identify complete products belonging to different primer sets in each sample. We plotted the percentage of expected products found in each sample with each primer set as a detection matrix for both positive samples and negative controls (Figure 4A). Because each value is a percentage of the 48 predicted sequences, 2.1% corresponds to at least one of the predicted complete products. With their own primer set, close to 100% of the 48 predicted sequences were found in almost all samples, whereas almost no complete sequences were retrieved with a different primer set.

**Figure 4.**
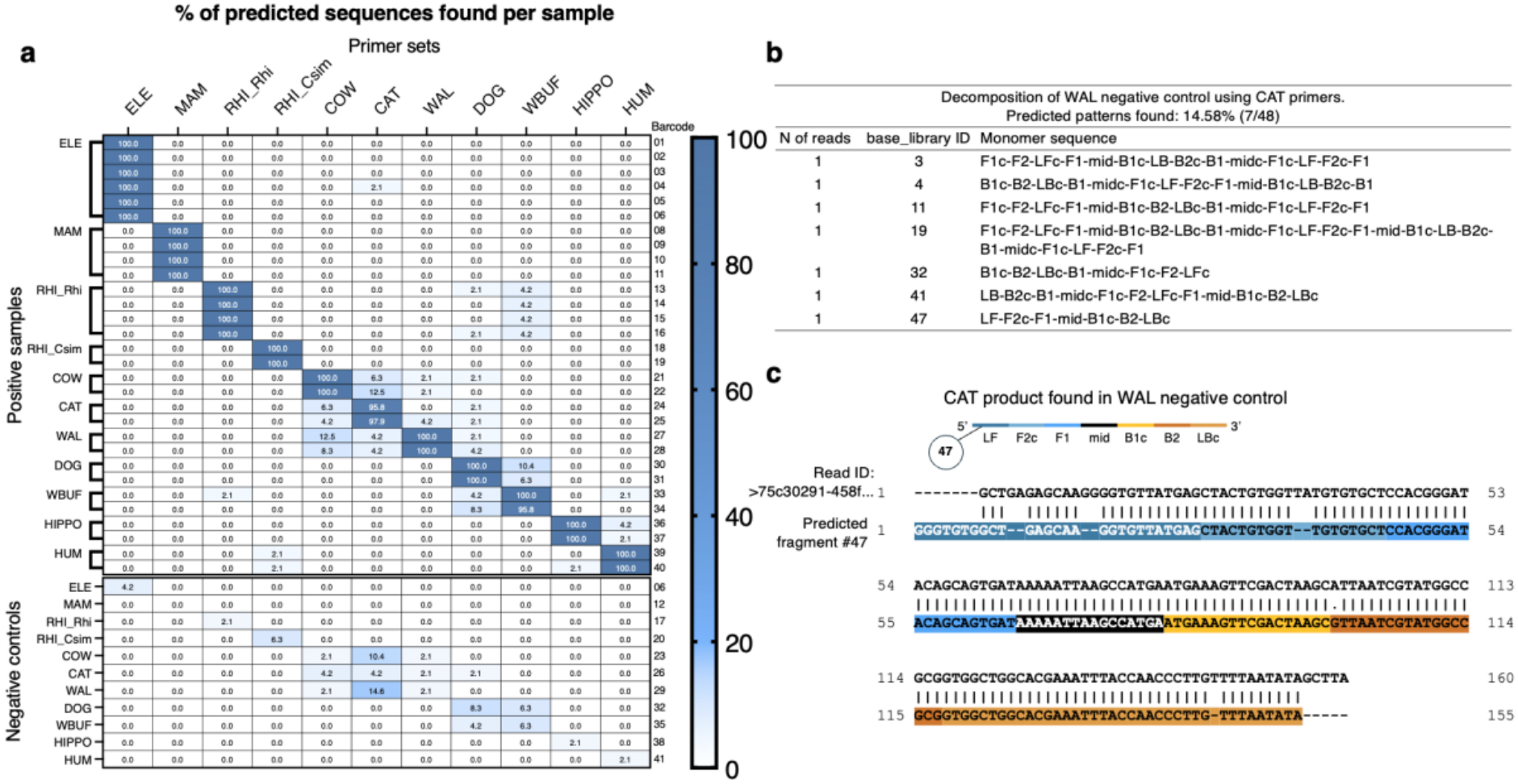
LAMPdecomposer has single read sensitivity. **a**. Detection matrix showing the percentage of the 48 predicted sequences found when each sample (rows) is decomposed with each primer set (columns). Positive samples are shown at the top and negative controls at the bottom, and each row is one sequencing reaction. The barcode labels match the original paper by Yugovich et al.^20^. A value of 2.1% corresponds to a single complete product. **b**. Decomposition of the WAL negative control with CAT primers. Seven reads were found, each containing a different complete CAT product (7 of 48 predicted sequences). The table gives the number of reads, the base library ID and the monomer sequence of each product. **c**. Alignment of a read from the WAL negative control (read ID 75c30291-458f…) with the predicted CAT fragment #47 (LF-F2c-F1-mid-B1c-B2-LBc). Coloured blocks mark monomer regions, vertical bars mark matching bases, dashes mark gaps and dots mark contradicting bases. The read contains the complete product at the monomer level.

Nevertheless, a few complete products from other primer sets were found in some positive samples, for example COW, CAT, and WAL products were found in each positive sample when samples were decomposed with any of the other two primer sets, and the same occurred to DOG and WBUF. These products correspond to between 1 and 8 reads per sample, accounting for up to 6 different predicted reads. Negative controls also contained a few products from their own primer set, between 2.1% and 8.3% of the predicted sequences (Figure 4A). The majority of reads decomposed in negative sequences showed a disorganised primer sequence, at very low prevalences, indicating those reads were likely primer dimers (Supplementary Table 1).

Complete products from other primer sets were also retrieved from some negative controls. The most prominent example is the WAL negative control decomposed with CAT primers. In this control, LAMPdecomposer identified 7 reads out of 2113 in the sample, each containing a different complete CAT product, the longest of which was 24 monomers long (Figure 4B). Alignment of the read containing product #47 confirms that the expected CAT sequence is complete at the monomer level in the WAL negative control (Figure 4C). In either case, LAMPdecomposer successfully decomposed targets to single read sensitivity.

### Relative prevalence of products shows a preferred path for LAMP products

Having established that the predicted products are consistently detected across different primer sets, we next investigated whether their relative abundances could provide insights into the mechanisms of LAMP amplification. We hypothesised that, although multiple pathways can generate the same products, differences in their prevalence might reveal preferential routes of product formation. To investigate this, we mapped the mean prevalence of each predicted sequence onto the amplification product tree (Figure 5).

**Figure 5.**
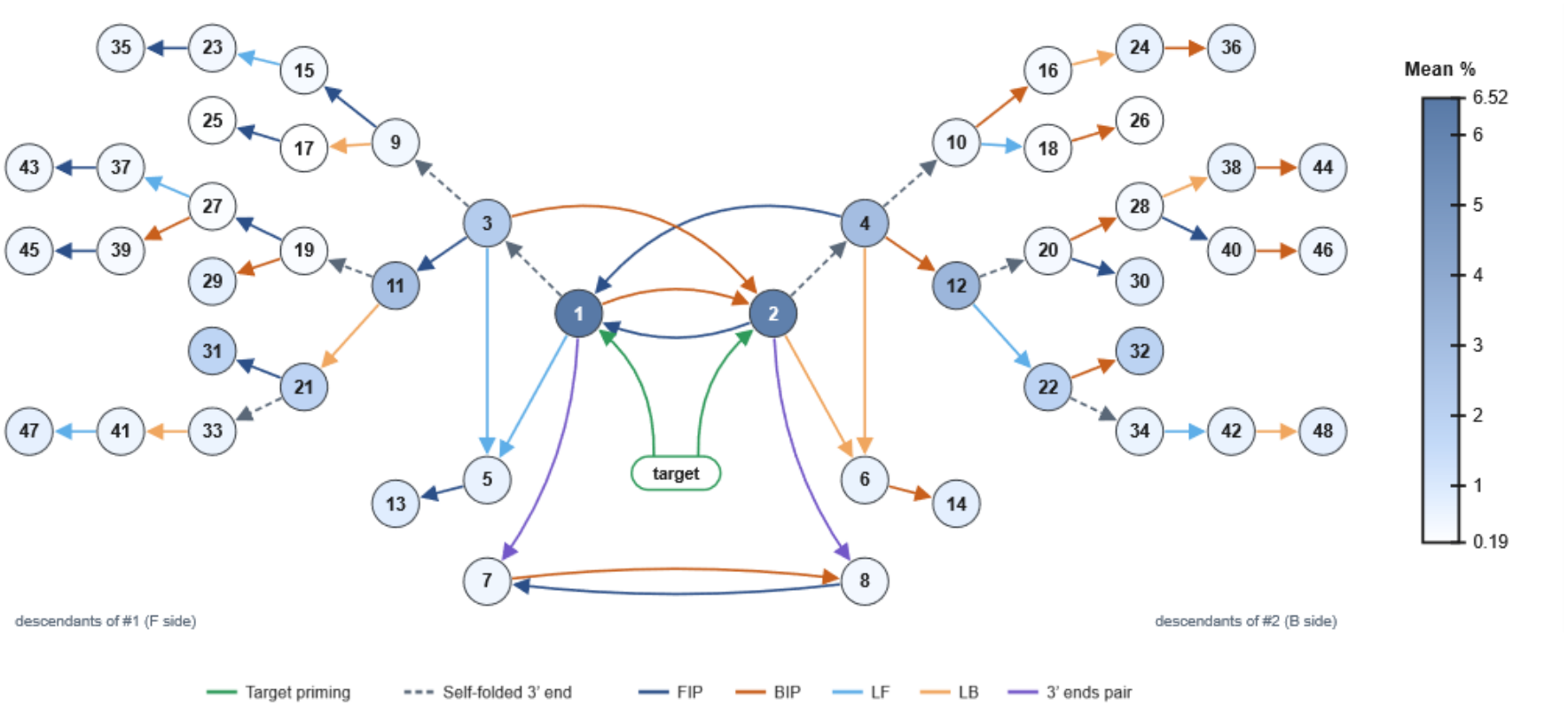
Relative prevalence of LAMP products reveals a preferred amplification path for the formation of amplification products. Mean prevalence of each base library sequence mapped onto the amplification product tree from Figure 2B. Node colour indicates mean prevalence as a percentage of all amplification reads across the 30 positive samples from 11 primer sets, ranging from 6.52% to 0.19%. The first amplification seeds, #1 and #2, are the most prevalent sequences. Prevalence remains high along the preferred path through #3/#4, #11/#12 and #21/#22, drops at intermediate sequences on all other branches and rises again at the extremities. Arrow colours and layout are as in Figure 2B.

Analysis of the prevalence of amplification products across all samples and primer sets shows that the most common sequences are the first amplification seeds, #1 and #2, with prevalences of 6.52% and 6.19% of the size filtered amplification reads, respectively (Figure 5). The most notable result is that sequences further down the tree are usually more common than their parents. This is the case for the early sequences #11, #12, which are more prevalent than #3 and #4, and for #13, #14, #23, #24, #25, #29, #30, #32, #35, #37, #38, #39, #40, #42, #43, #45, #46, #47 and #48. The only exceptions are #26, #31, #36 and #44, which are less prevalent than their parents by only 0.03, 0.01, 0.01 and 0.02 percentage points, respectively. The relative prevalences also reveal a preferred path of product formation, running through #3/#4, #11/#12 and #21/#22. These sequences are more prevalent than any of their other child sequences. Along all other branches, prevalence drops at intermediate sequences and rises again at the extremities.

Together, these findings reveal reproducible differences in the relative abundance of predicted LAMP products and identify a putative preferential amplification pathway. Although product prevalence does not directly establish the kinetics or directionality of individual reactions, mapping these abundances onto the predicted product tree provides insights into how different amplification mechanisms may contribute to the final product mixture. This approach therefore extends the utility of LAMPdecomposer beyond product identification, enabling sequence-level investigation of the pathways underlying LAMP amplification. An animation of the sequences from the highest mean prevalence to the lowest can be found in Supplementary File 2.

## Discussion

Here, we present LAMPdecomposer, a computational framework that combines a predictive model of loop-mediated isothermal amplification (LAMP) with nanopore sequencing to resolve the sequence diversity of concatemeric amplification products. We established a generalisable library of 48 predicted sequences generated through four molecular mechanisms and demonstrated that nearly all predicted products were present across 11 different primer sets. Despite the structural complexity of LAMP products, this relatively small library explained 52.2% of sequencing reads within its defined length range, while expanding the library to 600 sequences yielded comparatively modest improvements in coverage. This long tail is consistent with the proposed mechanism of product formation. The longest reads are likely those that the polymerase had the most time to extend, that is, those that began extending earliest in the reaction. Because the number of ways a sequence can fold and be extended grows with its length, products containing many monomers are expected to be largely unique, and ever-larger sample sizes would be needed to observe the same one twice. Therefore, expanding the library yields diminishing returns, as each additional sequence accounts for a progressively smaller fraction of the reads.

Mapping the relative abundance of individual products onto the predicted amplification tree also revealed non-uniform product distributions and a putative preferential amplification pathway. These findings suggest that much of the observed complexity of LAMP amplification can be described by a limited set of recurring molecular structures and mechanisms. Importantly, because the model is defined by the arrangement of primer-binding regions rather than target-specific nucleotide sequences, it can be adapted to different LAMP assays, providing a general framework for investigating amplification products at the sequence level.

Beyond characterising product diversity, LAMPdecomposer provides a means of assessing amplification specificity and identifying potential contamination. By decomposing nanopore reads into ordered patterns of primer target regions, the method enables the identification of complete amplification products originating from different primer sets, even when represented by a single sequencing read. We demonstrated this capability by identifying products associated with other primer sets in both positive samples and negative controls, including seven reads corresponding to complete products from a different target in one negative control. This is particularly relevant to molecular diagnostics, where the high amplification efficiency of LAMP increases the risk of amplicon contamination and false-positive results. Unlike approaches that assess amplification through fluorescence or product size alone, LAMPdecomposer provides sequence-level evidence of product identity, allowing unexpected amplification products to be distinguished from those predicted for the intended target. The consistent recovery of the predicted sequences across primer sets further suggests that the base library could provide a reference for assessing whether a reaction generates the expected amplification products. Although LAMPdecomposer alone cannot distinguish closely related targets amplified by the same primer set, SNPdetector resolves them in a post-processing step when a degenerate primer is used and the target sequences are supplied. In the dataset we evaluated, it assigned all ten samples to one of the tested species. Together, these capabilities offer a complementary approach to conventional assay validation and quality control, particularly for diagnostic applications requiring high specificity.

Several limitations remain. Although nearly all predicted sequences in the library were detected in every positive sample across 11 different primer sets, a substantial proportion of sequencing reads could not be explained by the library. Although most of these unexplained reads contained more monomers than the longest predicted sequences (suggesting longer concatemers not covered by our model), others may represent alternative amplification products or structures arising through mechanisms not represented in the library. Although each unexplained pattern was individually rare, together they accounted for a considerable fraction of the reads. Furthermore, nanopore sequencing provides a snapshot of the accumulated amplification products rather than a direct measurement of reaction kinetics. Consequently, the preferential pathways inferred from product abundance remain hypotheses that could be investigated through time-resolved sequencing experiments. Future developments could extend the model to longer and less common products, incorporate additional amplification pathways and evaluate its performance across a wider range of targets and experimental conditions. By providing a generalisable method for resolving the molecular composition of LAMP products, LAMPdecomposer establishes a foundation for mechanistic studies of isothermal amplification and the development of more reliable molecular diagnostic assays.

## Methods

### Source of sequencing data, targets and primer sequences

Raw Oxford Nanopore reads were obtained from the test dataset of Yugovich et al. (2026)^20^, available at https://github.com/oliviayugovich/LAMPSEQ/tree/main/test_data/. The files are numbered sequentially from barcode01.fastq to barcode41.fastq, and the last barcode of each primer set is a sequenced negative control. The primer sequences, and the accession numbers of the sequences used to design them, were published in the same study^20^.

### Creation of the base library for concatemeric sequences

The base library of concatemeric sequences was built from a generalised structure containing the minimum set of regions present in any LAMP amplification seed, plus a mid region between F1 and B1c. This structure consists of two complementary strands: 5′-F1c-F2-LFc-F1-mid-B1c-LB-B2c-B1-3′ and 5′-B1c-B2-LBc-B1-midc-F1c-LF-F2c-F1-3′. The amplification products arising from each strand were predicted for every primer-binding region and grouped into four types: (i) products that become blunt-ended and double-stranded; (ii) products that release single strands back into the reaction (recycled); (iii) products with a self-priming 3′ end that can be extended into longer concatemers; and (iv) products formed by the interaction of two strands with complementary 3′ ends. Products containing the F3 and B3 regions were not considered, because they do not take part in the exponential phase of amplification.

The resulting tree (Figure 2b) mainly shows the new products created by each primer–target or self-folding interaction, because many binding events regenerate molecules already shown. Each of these events produces two strands: a new sequence and a regenerated copy of a strand already described. In such cases, only the new strand is added to the tree. Products formed by the annealing of the 3′ ends of two strands are not shown, except for sequences #7 and #8, which give rise to new products. The tree is therefore not an exhaustive map of all LAMP products. All amplification paths were mapped for the sequences closest to the centre of the tree, but each strand, particularly those towards the tips, can give rise to further, longer and more complex concatemers. These were not described, as they fall outside the scope of this work.

### Resolving concatemeric sequences with LAMPdecomposer

LAMPdecomposer (v1.0.0) requires three inputs: the LAMP primer sequences (FASTA), the target sequence the primers were designed against (FASTA), and the Oxford Nanopore reads (FASTA or FASTQ, optionally gzip-compressed). For each primer set, LAMPdecomposer locates every primer site on the target and generates a custom set of monomers (monomers.fa) and a library of expected amplicon patterns (library.tsv), following the rules described in Supplementary Note 1. Reads are then decomposed into these monomers with StringDecomposer v1.1.2^18^. LAMPdecomposer converts each read’s decomposition into a monomer pattern, that is, the ordered series of monomer hits along the read with reverse-complement hits marked, and compares it with library.tsv. A read is counted as explained when its pattern exactly matches a pattern in library.tsv. For each sample, LAMPdecomposer reports the proportion of explained reads, the library patterns detected, and tables of explained and unexplained patterns. It also writes a summary table covering all samples.

LAMPdecomposer is run with “python LAMPdecomposer.py --primers [primers.fasta] --target [target.fasta] --input [reads file or folder]”. The primer file contains the sequences of primers F3, B3, F2, F1c, B1c and B2 (required) and LF and LB (optional), written 5′→3′, and degenerate IUPAC bases are allowed. Full FIP and BIP sequences may be given instead of F1c/F2 and B1c/B2. In that case, the longest 3′ stretch matching the target (≥10 nt) is taken as F2 or B2, and the remaining 5′ part as F1c or B1c. The target file contains a single sequence spanning at least F3 to B3c. A longer sequence, such as the full sequence used for primer design, can also be used, and the region outside F3–B3c is ignored. The --input option accepts one or more read files and/or a folder, and each file is analysed separately. With the --filter-length option, reads that cannot be explained by the library because they contain fewer monomers than the shortest library pattern or more than the longest are excluded from the analysis. Primers were located on the target allowing up to two mismatches, and stretches of ≤10 nt between adjacent primer sites were merged into the neighbouring primer monomer (default settings). Of the StringDecomposer outputs, only final_decomposition.tsv was kept, and the files final_decomposition_alt.tsv and final_decomposition_raw.tsv are deleted to save space.

### Evaluation of base library size

To evaluate whether a larger base library would explain more reads, base libraries of 30, 48 and 600 sequences were compared. Reads were first decomposed with the 48-sequence base library. The read counts of all positive samples were then pooled by read size (in monomers) and split into reads explained and reads not explained by the library. The 30 most prevalent sequences of this library formed the 30-sequence library. To build the 600-sequence library, the read patterns in the positive samples of primer sets ELE and MAM were ranked by prevalence. For each primer set, patterns that were also present in the negative control were removed, as they were likely to be primer dimers. The remaining patterns were converted back into base-library form, and the most prevalent patterns from both primer sets were combined into a 600-sequence library that also retained the 48 original sequences. Because this library was trained on ELE and MAM, it was tested only on the other primer sets. The 600-sequence library was built with LampTrainLibrary.py using the command “python LampTrainLibrary.py results --set ELE:1-6:7 --set MAM:8-11:12 --size 600”, and the three libraries were compared with LampLibrarySizes.py using the command “python LampLibrarySizes.py results --library 30 --library 48 --library results/base_library_600.txt”.

### Species assignment with SNPdetector

When a primer set targets more than one species, primers can carry degenerate nucleotides (IUPAC codes) at each position where the target species differ. A degenerate primer is synthesised as a mixture of every possible sequence version. The version that perfectly matches the template anneals and is extended more efficiently than mismatched versions, so it becomes enriched in the amplicons. SNPdetector expands a degenerate primer into all of its versions and counts the reads that contain each version exactly, both as written (forward) and as its reverse complement (reverse). The user gives each target sequence a name, for example the species, and the primer version that matches that target’s binding site is reported under that name. Because LAMP concatemers contain alternating sense and antisense copies of the target, the forward and reverse counts of a version should be similar, and they are treated as two measurements of that version. The most abundant version (highest mean of forward and reverse counts) is compared with every other version by a one-sided two-sample Student’s t-test (equal variances, degrees of freedom = 2). A sample is assigned to the species named on the leading version only when that version is significantly more abundant than all others (p < 0.05 by default). Otherwise, the sample is reported as inconclusive.

SNPdetector is run from the terminal (python SNPdetector.py). It asks for the primer name and sequence, then for one or more target names and sequences, and then for the read files or a folder containing them. Each target can be typed as a sequence or given as a FASTA file. All inputs can also be given as command-line arguments. How primer versions are generated, and recommendations for designing LAMP assays to discriminate SNPs, are described in Supplementary Note 2.

## Data and code availability

The raw nanopore reads analysed in this study were generated by Yugovich et al.^20^ and are available in the LAMPSEQ repository (https://github.com/oliviayugovich/LAMPSEQ,test_data/00_raw_data, commit 2823315). Target sequences were obtained from NCBI using the accessions listed in the Supplementary material of the same study. LAMPdecomposer (v1.0.0) and the scripts used to generate the data for all figures and tables are freely available under the GNU GPL v3.0 at https://github.com/Carlos-dosSantos/LAMPdecomposer and archived at Zenodo (https://doi.org/10.5281/zenodo.24539001).

## AI disclosure

Generative AI (Claude Opus 5.5, Anthropic, “High” mode, October 2026) was used under the constant supervision of the lead author (C.A.S.) to write the Python scripts presented in this study. Claude was also used to write the HTML code for the animations of the base library tree and prevalence in Supplementary Files 1 and 2, and Figures 2b and 5 show static snapshots of these animations. Each animated step was based on a static figure created by C.A.S. in PowerPoint, which showed the binding steps needed to create the products shown in the tree. The authors take full responsibility for the content of this paper.

## Acknowledgements

This work was supported by a Royal Society Dorothy Hodgkin Research Fellowship DKR00620 and a BBSRC project grant UKRI3595 (to N.C.R.), and an Institute for Global Pandemic Planning–funded PhD at the University of Warwick, UK (to C.A.S).

## Contributions

C.A.S conceived the study and performed the primary analyses. A.M. wrote code and provided analytical guidance. N.C.R. provided supervision and funding. C.A.S wrote the initial manuscript draft with assistance from N.C.R. All authors reviewed and approved the final manuscript.

## Competing interests

The authors declare no competing interests.

## Notes

https://github.com/Carlos-dosSantos/LAMPdecomposer

https://doi.org/10.5281/zenodo.24539001

### Competing Interest Statement

The authors have declared no competing interest.

