## Supplementary material for "LAMPdecomposer: A generalisable framework for predicting and identifying loop-mediated isothermal amplification products using nanopore sequencing"

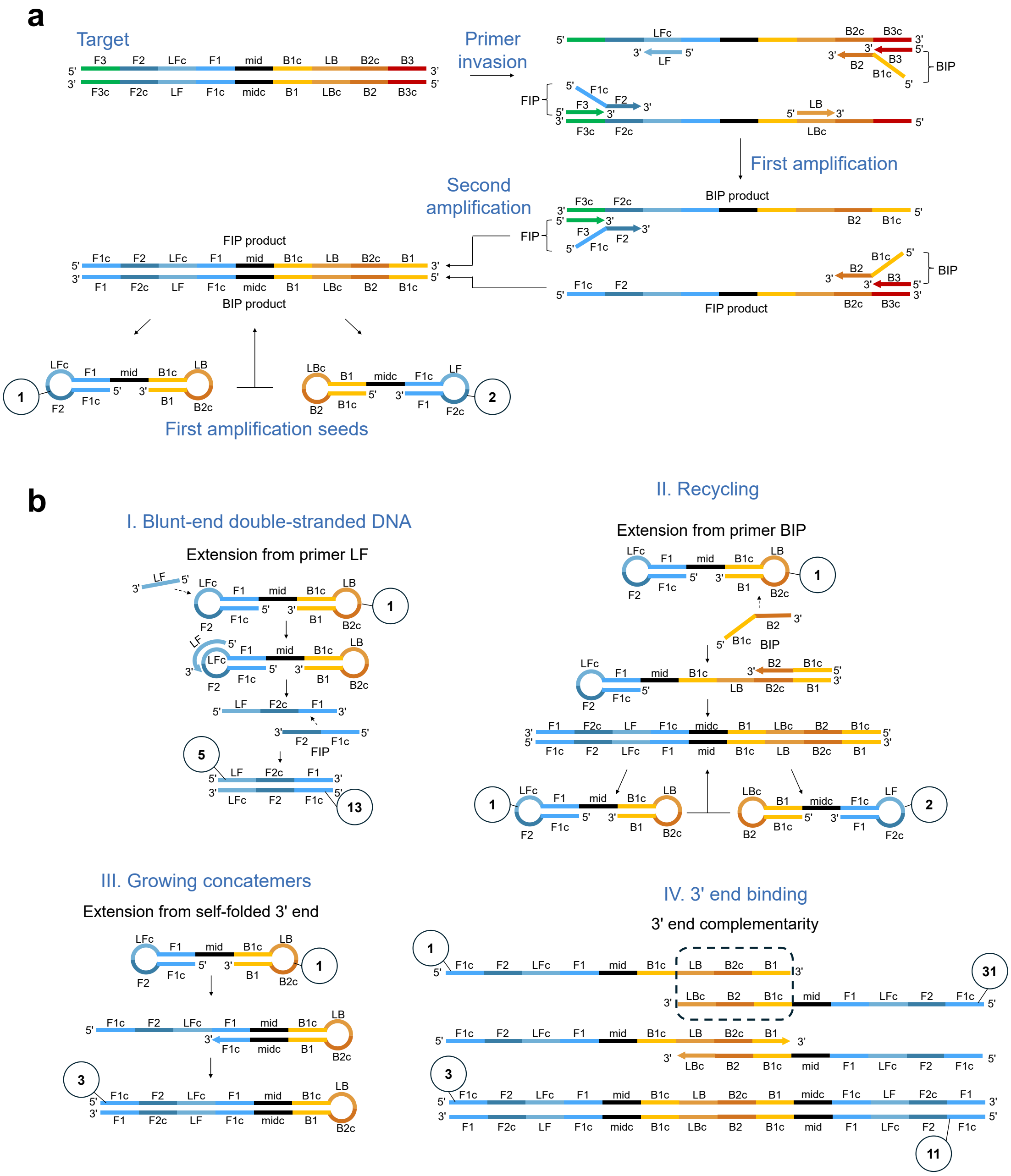

***Supplementary Figure 1. Mechanisms of LAMP product formation. a.*** *Generation of the first amplification seeds.* ***b.*** *The four mechanisms that explain all products in the base library. As in Figure 1, however specific sequences are numbered according to their assignments in the amplification product tree.*

**Supplementary Table 1. *Decomposition results for the 11 negative controls in each primer set.*** *The most common patterns are not predicted by the library and are the result of primer dimers.*

| Datasets | Primer set | Total reads (raw file) | Total reads (size filtered) | Reads explained by library | % filtered reads explained | average % of predicted sequences found | Most common pattern (5' - 3') |
|---|---|---|---|---|---|---|---|
| 1 | ELE | 1,213 | 953 | 2 | 0.21% | 4.17% | B1c-B2-B1c-LFc-B2c-B1 |
| 1 | MAM | 328 | 193 | 0 | 0.00% | 0.00% | B1c-B2-F1c-F2c-F1 |
| 1 | RHI_Rhi | 10,347 | 7,632 | 1 | 0.01% | 2.08% | F1c-F2-B1c-B2c-B1 |
| 1 | RHI_Csim | 569 | 364 | 3 | 0.82% | 6.25% | B1c-B2-F2-B1 |
| 1 | COW | 1,642 | 1,126 | 5 | 0.44% | 2.08% | F1c-F2-F1 |
| 1 | CAT | 648 | 384 | 3 | 0.78% | 4.17% | B1c-B2-B2c-B1 |
| 1 | WAL | 2,113 | 1,650 | 1 | 0.06% | 2.08% | B1c-B2-F3-B3c-B2c-B1 |
| 1 | DOG | 1,414 | 976 | 9 | 0.92% | 8.33% | B1c-B2-F1c-B3c-LFc-LFc-B3c-LBc |
| 1 | WBUF | 344 | 213 | 3 | 1.41% | 6.25% | mid-B1-midc-B1c |
| 1 | HIPPO | 1,135 | 821 | 1 | 0.12% | 2.08% | B1c-B2-F3-B3c-B2c-B1-B1c-B2-F3-B3c-B2c-B1 |
| 1 | HUM | 183 | 106 | 1 | 0.94% | 2.08% | B1c-B2-B2c-B1-B1c-B2-B2c-B1-B1c-B2-B2c-B1 |

**Supplementary Note 1.** Translating a LAMP target into monomers

To resolve LAMP concatemers with StringDecomposer[1], the complete target region must first be translated into a set of monomers. A standard LAMP assay uses six primers (F3, B3, FIP, BIP, LF and LB), of which the loop primers LF and LB are optional. Because FIP and BIP each span two binding regions, the six primers bind eight regions, which on the leading strand (5′→3′) are F3, F2, LFc, F1, B1c, LB, B2c and B3c (Supplementary Figure 2a).

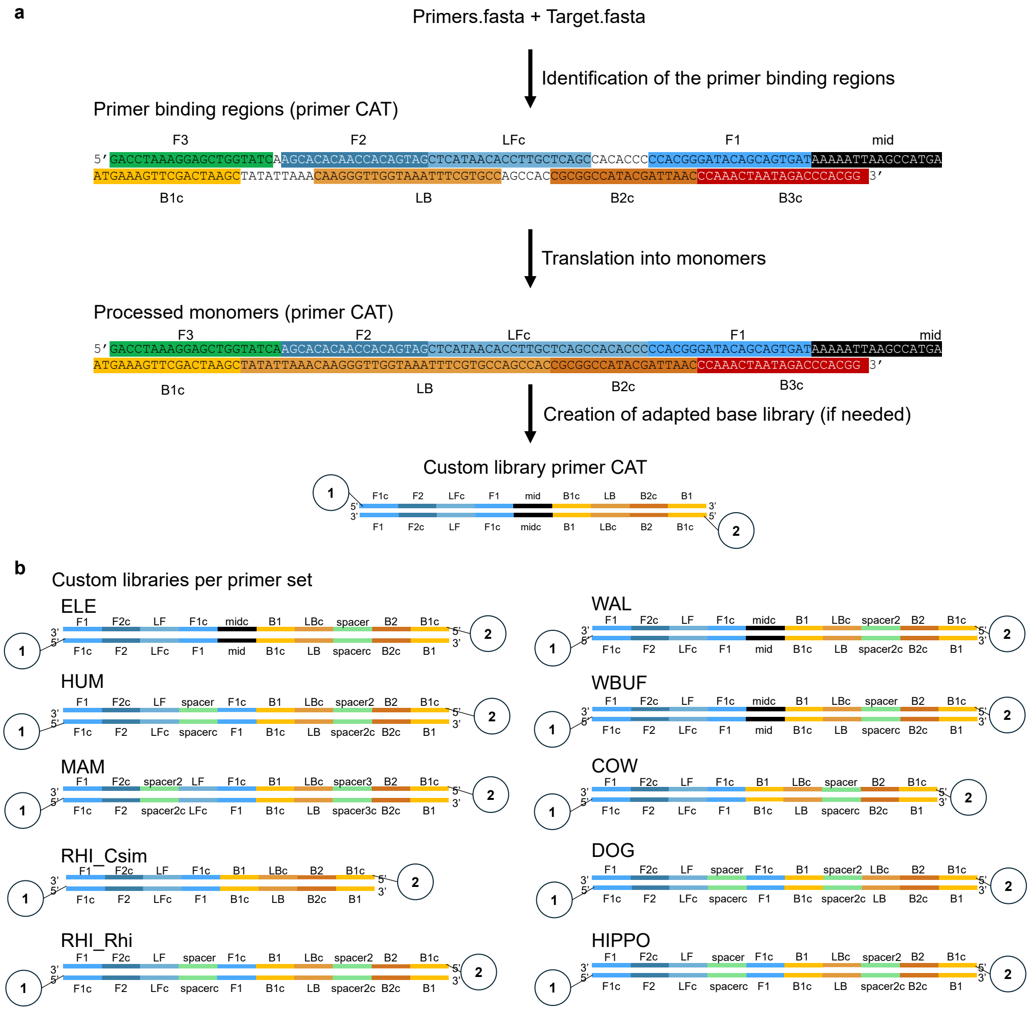

***Supplementary Figure 2. Translation of a LAMP target into monomers and adaptation of the base library. a.*** *Target sequence of the CAT primer set, shown on the forward strand. When the primers and the target region are supplied, LAMPdecomposer identifies the primer binding regions (top sequence), and then translates those regions into monomers (bottom sequence). In the monomer view, inter-primer sequences of 10 nucleotides or fewer have been added to neighbouring monomers, and longer ones have become monomers of their own, as with mid in CAT. The monomers are then used to adapt the base library when the primer set has named inter-primer monomers or lacks a loop primer.* ***b****. Dumbbell structures 1 and 2 of the base library, adapted for CAT and for each of the other ten primer sets (ELE, HUM, MAM, RHI_Csim, RHI_Rhi, WAL, WBUF, COW, DOG and HIPPO). Each shows the order of monomers on both strands, including the mid and spacer monomers specific to that primer set. Structures are numbered as in the base library. Raw data obtained from Yugovich et al. 2026.*[2]

The monomer set must cover the entire target region, including the sequences between primer binding regions. Inter-primer sequences of 10 nucleotides or fewer are added to a neighbouring monomer according to the rules in Supplementary Table 2. F2, B1c and B2c are never extended, so that no sequence is added at the junctions within FIP (F1c–F2) and BIP (B1c–B2). Inter-primer sequences longer than 10 nucleotides become monomers of their own. The sequence between F1 and B1c is named “mid”, and the others are named “spacer”, “spacer2”, “spacer3” and so on, in 5′→3′ order (Supplementary Figure 2a). Degenerate primers are expanded into all of their sequence versions, and each version becomes a separate monomer.

The monomer set is then used to adapt the base library of expected LAMP product structures to the primer set. Named inter-primer monomers are inserted wherever their two flanking regions are adjacent, and the region of any absent loop primer is removed (Supplementary Figure 2b). The monomer sequences and the sequencing reads are given to StringDecomposer, which decomposes each read into an ordered series of monomers. A read is considered explained when its monomer series exactly matches a pattern in the adapted library, and the percentage of library patterns found in the reads is reported.

***Supplementary Table 2.*** *Assignment of inter-primer sequences of 10 nucleotides or fewer to neighbouring monomers. Positions are given on the leading strand (5′→3′). The last two rows apply only to assays without loop primers.*

| Inter-primer sequence between | Added to |
|---|---|
| F3 and F2 | 3’ end of F3 |
| F2 and LFc | 5’ end of LFc |
| LFc and F1 | 3’ end of LFc |
| F1 and B1c | 3’ end of F1 |
| B1c and LB | 5’ end of LB |
| LB and B2c | 3’ end of LB |
| B2c and B3c | 5’ end of B3c |
| F2 and F1 (no LF primer) | 3’ end of F2 |
| B1c and B2c (no LB primer) | Always its own monomer |

**Supplementary Note 2.** Degenerate primer versions and species assignment with SNPdetector

A degenerate primer carries an IUPAC ambiguity code at one or more positions. Each code represents two or more possible nucleotides; for example, Y represents C or T, and N represents any nucleotide. During synthesis, every nucleotide represented at a degenerate position is incorporated at approximately equal frequency, so the ordered primer is a mixture of every possible sequence version. The number of versions is the product of the number of nucleotides represented at each degenerate position: one two-base code gives two versions, and each additional two-base code doubles that number. The ELE LB primer (5′-AAYCGATTYTTCGCCYTCC-3′) contains three Y positions and therefore consists of $2^3 = 8$ versions, each present at about one-eighth of the total LB concentration. All eight versions take part in the amplification. However, the primer was designed to discriminate only three species (Elephas maximus, Loxodonta africana and Loxodonta cyclotis), so five of the versions match the targets partially.

A primer version that perfectly matches the template anneals and is extended more efficiently than mismatched versions, so it is preferentially incorporated into the amplification products. Versions with one mismatch are expected to be incorporated more often than versions with two or three. The effect of a mismatch also depends on its position, and mismatches near the 3′ end are the most disruptive. In a positive sample, the version that matches the template should therefore be the most abundant version in the reads.

SNPdetector expands a degenerate primer into all of its versions and counts the reads that contain each version and its reverse complement. The user gives each target sequence a name, and the version that matches that target is reported under that name. The versions that match no target show how often mismatched primers are incorporated. SNPdetector then tests whether the most abundant version is significantly more abundant than every other version (Methods). If it is, the sample is assigned to the species named on that version. The number of reads containing each LB version is shown in Supplementary Figure 3 for barcodes 01–07 (ELE primer set) and 13–17 (RHI_Rhi primer set).

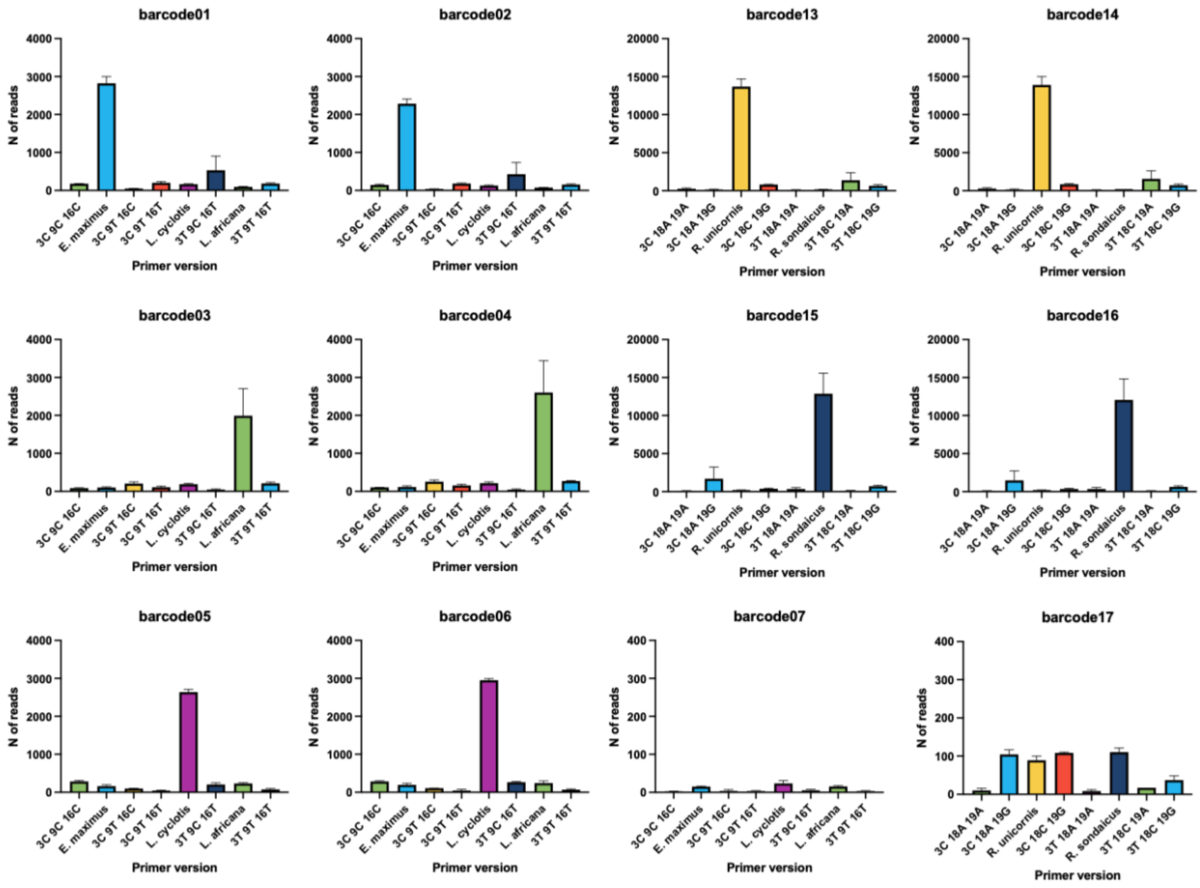

***Supplementary Figure 3. Read counts for each version of the degenerate LB primers.*** *SNPdetector counted the reads containing each version of the LB primer in the ELE primer set (barcodes 01–07) and the RHI_Rhi primer set (barcodes 13–17). Each LB primer has three degenerate positions and therefore eight versions. Versions are labelled by the nucleotides at those positions: 3, 9 and 16 for ELE LB, and 3, 18 and 19 for RHI_Rhi LB. A version that matches a target species is labelled with that species' name instead. Bars show the mean of the forward count (reads containing the version as written) and the reverse count (reads containing its reverse complement). Error bars show the standard deviation of these two counts. The samples are* Elephas maximus *(barcodes 01–02),* Loxodonta africana *(03–04),* Loxodonta cyclotis *(05–06),* Rhinoceros unicornis *(13–14) and* Rhinoceros sondaicus *(15–16). Barcodes 07 and 17 are the no-template controls for ELE and RHI_Rhi, respectively. The y-axis scale differs between panels for ELE and RHI_Rhi in the no-template controls, which show a 10x smaller scale. Species assignments and p values are given in Figure 3e. Raw data from Yugovich et al., 2026.*[2]
